# Reconstruction of body mass evolution in the Cetartiodactyla and mammals using phylogenomic data

**DOI:** 10.1101/139147

**Authors:** E. Figuet, M. Ballenghien, N. Lartillot, N. Galtier

## Abstract

This preprint has been reviewed and recommended by Peer Community In Evolutionary Biology (http://dx.doi.org/10.24072/pci.evolbiol.100042).

Reconstructing ancestral characters on a phylogeny is an arduous task because the observed states at the tips of the tree correspond to a single realization of the underlying evolutionary process. Recently, it was proposed that ancestral traits can be indirectly estimated with the help of molecular data, based on the fact that life history traits influence substitution rates. Here we challenge these new approaches in the Cetartiodactyla, a clade of large mammals which, according to paleontology, derive from small ancestors. Analysing transcriptome data in 41 species, of which 22 were newly sequenced, we provide a dated phylogeny of the Cetartiodactyla and report a significant effect of body mass on the overall substitution rate, the synonymous vs. non-synonymous substitution rate and the dynamics of GC-content. Our molecular comparative analysis points toward relatively small Cetartiodactyla ancestors, in agreement with the fossil record, even though our data set almost exclusively consists of large species. This analysis demonstrates the potential of phylogenomic methods for ancestral trait reconstruction and gives credit to recent suggestions that the ancestor to placental mammals was a relatively large and long-lived animal.

## INTRODUCTION

Accessing the ancestral states of a trait is of primary importance to study the processes having shaped its evolution (Donoghue 1989; Cunningham et al. 1998). Yet in most cases, ancient species and their characteristics are no longer observable and can only be indirectly estimated. In some cases, information on ancestral morphology or anatomy can be obtained through paleontological data. For instance, the analysis of a large fossil data set from marine environments was consistent with so-called Cope’s or Depéret’s rule, which postulates a tendency for lineages to evolve towards larger body mass (Heim et al. 2015; Bokma et al. 2016). Fossils, however, are scarce and their interpretation is often controversial, so that paleontology can but rarely provide us with a complete and detailed picture of the evolution of a trait in a clade.

To fill the gaps of direct evidence, evolutionary biologists have for long turned their attention onto the development of estimation methods using the comparative approach. This is typically achieved by fitting a model of trait evolution on a phylogeny to the observed data at the tips of the tree. For continuous character, such as body mass, the most basic model assumes an unidimensional Brownian motion with constant rate across branches (Felsenstein 1985). Since then, more complex models have been proposed, such as models assuming an Ornstein-Uhlenbeck process, which constantly pulls the trait toward an optimum value (Beaulieu et al. 2012), or models seeking for a preferential trajectory in the random walk (Hunt 2006). The major impediment of these approaches is that they rely on a unique realization of the modelled process. Consequently, they lack statistical power and are sensitive to model misspecification (Oakley and Cunningham 2000).

The quickly accumulating molecular data might convey a new source of information about ancestral traits. Molecular evolution is influenced by a number of factors related to species biology and ecology, such as mutation rate, effective population size and mating systems (Lynch 2007), which in turn are presumably influenced by species life-history traits. Genomic sequences are expected to keep track of the various selective or neutral pressures that species have undergone through time. The idea, therefore, would be to indirectly estimate ancestral traits based on ancestral molecular reconstructions. The strength of this approach comes from the fact that molecular characters are numerous, which considerably increases the power and accuracy of phylogenetic inference methods.

In mammals, various aspects of the molecular evolutionary process have been linked to species life-history traits (LHT), including nucleotide and amino acid substitution rates (Martin and Palumbi 1993; Bromham et al. 1996; Nabholz et al. 2008) and genomic GC content (Romiguier et al. 2013; Lartillot 2013). The nonsynonymous (= amino acid changing, dN) to synonymous (dS) substitution rate ratio, dN/dS, is of specific interest, as it has been found to be positively correlated with species body size, longevity and generation time in mammals (Nikolaev et al. 2007; Popadin et al. 2007; Romiguier et al. 2012). These relationships were interpreted as reflecting a decreased effective population size (Ne) in large mammals (Damuth 1981, 1987; White et al. 2007), compared to small-sized ones. According to the nearly neutral theory of molecular evolution (Ohta 1987), purifying selection against slightly deleterious mutations is less efficient in small populations, in which the random effects of genetic drift dominate. Assuming that a majority of nonsynonymous changes are deleterious, an increased proportion of nonsynonymous substitutions is therefore expected in (large-bodied) low-Ne species (Figuet et al. 2016). Besides dN/dS, an increased effective population size in small-sized species is also expected to impact the efficiency of biased gene conversion (gBGC), a recombination-driven process that favors the spread of GC relatively to AT alleles, mimicking a selective advantage for G and C (Galtier et al. 2001; Duret and Galtier 2009; Webster et al. 2005). Consistent with theoretical predictions, it was shown that small-sized, large-Ne mammals tend to exhibit a high GC-content compared to large-sized species, particularly at third codon positions (Romiguier et al. 2010).

Recently, a few studies took advantage of these correlations between species life-history traits and molecular evolutionary processes to estimate ancestral character states in mammals using phylogenetic approaches. Using hundreds of nuclear genes, Romiguier et al. (2013) reported that the estimated dN/dS ratio and GC content dynamics during early divergence in placental mammals were comparable to those of current long-lived mammals, and very different from those of current short-lived mammals. They concluded that placental ancestors were probably long-lived (>20 years) animals, in sharp contrast with the classical view of tiny early mammals diversifying after the massive extinction of large reptiles (e.g. Feldhamer et al. 2007). Lartillot and Delsuc (2012) analysed a smaller data set using a more sophisticated method in which the correlation between molecular rates and LHT was explicitly modelled (Lartillot and Poujol 2011). Their results were largely consistent with those of (Romiguier et al. 2013), rejecting the hypothesis of mouse-like placental ancestors, even though point estimates for ancestral traits differed between the two studies.

These estimates of ancestral LHT, which concern early placental ancestors, can hardly be compared to paleontological data owing to the uncertainty regarding placental origins (Archibald et al. 2001; Asher et al. 2005). Molecular dating, for instance, places the placental root in the Cretaceous (dos Reis et al. 2012; Reis et al. 2014), but no fossils from this epoch (Goswami et al. 2011; Wible et al. 2007, 2009) are unambiguously assigned to Placentalia. Perhaps for this reason, the results of Lartillot and Delsuc (2012) and Romiguier et al. (2013), which rely on novel, untested methods, so far have had a limited impact on the paleontological literature. To assess the reliability of molecular-aided reconstructions of past life-history traits, one idea would be to focus on clades for which the fossil record provides a well-supported and consensual evolutionary scenario.

The clade of cetartiodactyls appears as an interesting candidate in this respect. Cetartiodactyls are a group of 332 recognized species, classified in 22 families (Hassanin et al. 2012), that emerged around the Cretaceous-Paleogene crisis according to molecular dating (Meredith et al. 2011; Psouni et al. 2012). It comprises, among others, clades such as suines (pig and peccaries), tylopodes (camels), true ruminants (giraffe, cow, antelopes) as well as hippos and cetaceans. Extant species of cetartiodactyls exhibit a wide spectrum of body mass, ranging from a few kilograms (mouse-deer, dwarf antelopes, pygmy hog) to tons (hippos, giraffes) and even hundreds of tons (whales). Furthermore, with the exception of cetaceans, they share a number of ecological similarities, such as a terrestrial life style and a herbivorous diet often associated with rumination, making their LHT perhaps more comparable than in Placentalia-scale analyses. The fossil record of Cetartiodactyla is quite furnished with a mean of six extinct genera per extant genus (McKenna and Bell 1997). Despite the majority of extant representatives of this clade exceeding a hundred of kilograms, the early fossil record consists in small-sized animals, such as the emblematic *Diacodexis*, the oldest known cetartiodactyl, which only weighted a few kilograms (Rose 1982). Besides *Diacodexis*, the Eocene record of cetartiodactyls, with families such as *Heloyidae, Raoellidae* which are close relative to whales or *Choeropotamidae* of which the genus *Amphirhagatherium*, only includes relatively small-sized species (Hooker and Thomas 2001; Orliac and Ducrocq 2011). It is therefore widely admitted that body mass in cetartiodactyls has been convergently increasing in a number of independent lineages during the evolution of this group.

This peculiar history of body mass evolution creates an engaging challenge for molecular data. Corroborating the existence of small-sized ancestral cetartiodactyls based on a sample of large-sized extant species would be quite reassuring regarding the reliability of these approaches. An analysis of complete mitochondrial genomes in 201 species of cetartiodactyls suggested that DNA-aided reconstruction can estimate ancestral LHTs more accurately than classical methods (Figuet et al. 2014). Mitochondrial DNA, however, is limited in size. In addition, it experiences a high rate of substitution and is thus potentially subject to mutational saturation, making ancestral reconstructions more challenging. Nuclear DNA, in contrast, shows a relatively low rate of evolution but is subject to recombination and GC-biased gene conversion. Although making the analysis potentially more complicated, these additional evolutionary forces also represent genuine sources of signal about ancestral traits through among-lineage variation in GC content.

In this study, we aim to evaluate the potential and robustness of genomic data for ancestral lifehistory trait reconstruction in the Cetartiodactyla. To this end, we generated a dataset of thousands of genes in 41 species, of which 22 were newly sequenced, and conducted analyses inspired by Romiguier et al. (2013) and Lartillot and Delsuc (2012). Firstly, we approached ancestral body mass, longevity and age of sexual maturity based on the observed correlation between life-history and molecular traits across extant species (Romiguier et al. 2013; Figuet et al. 2014). Secondly, we used the integrated approach developed by Lartillot and Poujol (2011), in which the correlated evolution of life-history traits and molecular parameters across the phylogenetic tree is modelled as a multivariate Brownian motion, so as to obtain quantitative estimates of ancestral body masses and credibility intervals. The strength of this approach is to account for phylogenetic inertia and various sources of uncertainty by combining all variables in a single Bayesian analysis.

## METHODS

### Sample collection, RNA extraction and sequencing

Twenty one Cetartiodactyla species were sampled and analyzed in this study, namely the Chacoan peccary, common warthog, hippo, pygmy hippo, okapi, giraffe, reindeer, tufted deer, fallow deer, red deer, sika deer, Siberian musk deer, nilgai, nyala, common eland, Kirk’s dik-dik, dama gazelle, Nile lechwe, bontebok, addax, and white oryx. One species of Perissodactyla, the white rhinoceros, was also newly sequenced here. All reads are available from the NCBI-SRA database, BioProject PRJNA388863. Coding sequences from twelve additional Cetartiodactyla species and seven outgroups (two Eulypotyphla, two Chiroptera, two Carnivora, and one Perrisodactyla) were retrieved from publicly available genomic or RNA-Seq data bases (see electronic supplementary material, appendix A, table S1).

### Orthologous genes and alignments

*De novo* transcriptome assembly was performed with a combination of the Abyss and Cap3 programs following strategy B in Cahais et al. (2012). The longest ORF of each predicted cDNA was kept, contigs not including any ORF longer than 200bp being discarded. Orthology was predicted using the OrthoMCL software (Li et al. 2003, see electronic supplementary material, appendix A for details). Predicted orthologous sequences were aligned using the MACSE program (Ranwez et al. 2011), which relies on amino acid translated sequences while allowing for frameshifts at nucleotide level. Obviously mis-aligned regions were semi-automatically cleaned out using home-made scripts. Briefly, this was done by scanning each sequence for small regions flanked by undetermined base pairs, and removing them if they shared less than 70% similarity with any other sequence. The 3,022 resulting alignments were restricted to positions having less than 30% missing data, returning a total length of 3.4Mb of coding sequences in 41 species, with 17% missing data. Because the Bayesian integrated method developed by Lartillot and Poujol (2011) can hardly handle a data set of that size, we created a *reduced* data set in which we only kept genes at which less than two Cetartiodactyla species missing. The criterion was satisfied by 1,223 alignments, leading to a concatenation of 1,5Mb with 12% missing data.

### Life-history traits

Life-history traits values for body mass, maximum longevity and age of sexual maturity were retrieved both from the AnAge database build 13 (de Magalhães and Costa 2009, male and female values for sexual maturity) and the PANTHERIA database (Jones et al. 2009). When several estimates per species were available we calculated the mean (see electronic supplementary material, table S1). All values were log_10_-transformed prior to perform any analyses. Maximum longevity and age of sexual maturity in *Pantholops hodgsonii* as well as age of sexual maturity in *Pteropus vampyrus*, which were missing from AnAge and PANTHERIA, were obtained from the Animal Diversity Web resource (http://animaldiversity.org/).

### Phylogenetic reconstruction, substitution mapping, branch-specific dN/dS

A phylogenetic tree was estimated using raxML version 8 (Stamatakis 2006) with a GTR+GAMMA model. We trusted the reconstructed phylogeny as far as within-Cetartiodactyl branching orders were concerned. Regarding the inter-ordinal relationships, which are controversial, we used two alternative topologies taken from the literature, i.e., (Eulipotyphla, ((Chiroptera, (Carnivora, Perissodactyla)), Cetartiodactyla)) (Nishihara et al. 2006) and (Eulipotyphla, ((Carnivora, Perissodactyla), (Chiroptera, Cetartiodactyla))) (Nery et al. 2012). We also considered two alternative branching orders for the Nile lechwe *Kobus* (see electronic supplementary material, appendix B). The substitution mapping procedure (Dutheil et al. 2012; Romiguier et al. 2012) was used to estimate the mean dN/dS in each branch of the uncalibrated tree across the 3,022 alignments, following Figuet et al. (2016, see electronic supplementary material, appendix A for details about the methodology). This approach is similar but faster than, e.g., full likelihood branch models such as implemented in codeml (Yang 2007).

### Clustering procedure

To assess the potential noise introduced by recent variation in effective population size or life-history traits in terminal branches that could impede their relation with long-term dN/dS estimated by substitution mapping, we also combined the signal from closely related terminal branches into a single data point. To achieve this, we defined monophyletic clusters of species and calculated one dN/dS ratio per cluster. Clusters were defined at the level of tribes in Ruminantia, at the family level in Suina and Tylopoda, and at the parvorder level in Cetacea. The dN/dS value of a cluster was obtained by summing the nonsynonymous and synonymous substitution counts across its terminal branches and calculating the ratio. Life-history trait values of a cluster were defined as the mean of the log10-transformed values of its representatives, either from our sampling or from all available species at this taxonomic level in the AnAge database build 13. This clustering procedure resulted in the creation of 19 clusters in Cetartiodactyla, of which eleven contained two or three species and eight contained a single species (see electronic supplementary material, table S2).

### Bayesian analysis of the correlated evolution between substitution rates and body mass

Alternatively, we used the integrated COEVOL method (version 1.5, Lartillot and Poujol 2011) to jointly assess the correlations between life-history traits, dS and dN/dS and estimate ancestral states, based on the *reduced* concatenated data set. The method models the correlated evolution of the analyzed variables by assuming a multivariate Brownian diffusion process parameterized by a covariance matrix, along a time tree constrained by fossil calibrations. All parameters are estimated in a Bayesian framework using Monte Carlo Markov Chains. The strength of the coupling between a molecular variable and a LHT was measured by the correlation coefficient (posterior mean estimate) between the variables, and the statistical support was evaluated based on the posterior probability (pp) of a positive (pp close to 1) or negative covariation (pp close to 0). dS and dN/dS were analyzed as dependent variables, either jointly or separately. Ten calibration points were used (see electronic supplementary material, table S3, appendix A for details).

### GC3 analysis

Ancestral GC-content at third codon positions (GC3) was estimated for all nodes of the tree separately for each of the 3,022 genes using the approach of (Galtier and Gouy 1998), implemented in the bpp_ML suite (Dutheil and Boussau 2008). This method uses a nonhomogeneous and nonstationary Markov model of nucleotide evolution to estimate branch-specific equilibrium GC-content in a maximum-likelihood framework on the uncalibrated tree. We then employed the procedure used in Romiguier et al. (2013) to correlate GC3 evolution to life-history traits. This means calculating, for each of eleven selected pairs of extant species (see electronic supplementary material, appendix A for details about the methodology), an index of GC3 conservation, defined as γ=-t/log(τ), where t is the divergence time, and τ is Kendall rank correlation coefficient of GC3 across genes between the two considered species. This index was also calculated for pairs of ancestral taxa and used for ancestral body mass estimation based on the correlation with body mass in extant pairs.

## RESULTS

### Phylogeny and molecular dating

We took benefit of our extensive genomic data set to produce a phylogeny of the Cetartiodactyla clade. Analysed with raxML, the reduced and complete data set supported topologies that were identical (figure 1) and in very good agreement with previously published phylogenies (Price et al. 2005; Agnarsson and May-Collado 2008; Hassanin et al. 2012). Of note, Tylopoda branch out as the sister group to all other Cetartiodactyla in this tree, contradicting the species-rich mitochondrial-based analysis of (Hassanin et al. 2012), in which Suina was the earliest diverging group. Bootstrap support was 100% for all nodes, with the exception of the position of the Nile lechwe, for which 88% and 78% bootstrap support were obtained from the complete and reduced data set, respectively.

**Figure 1.**
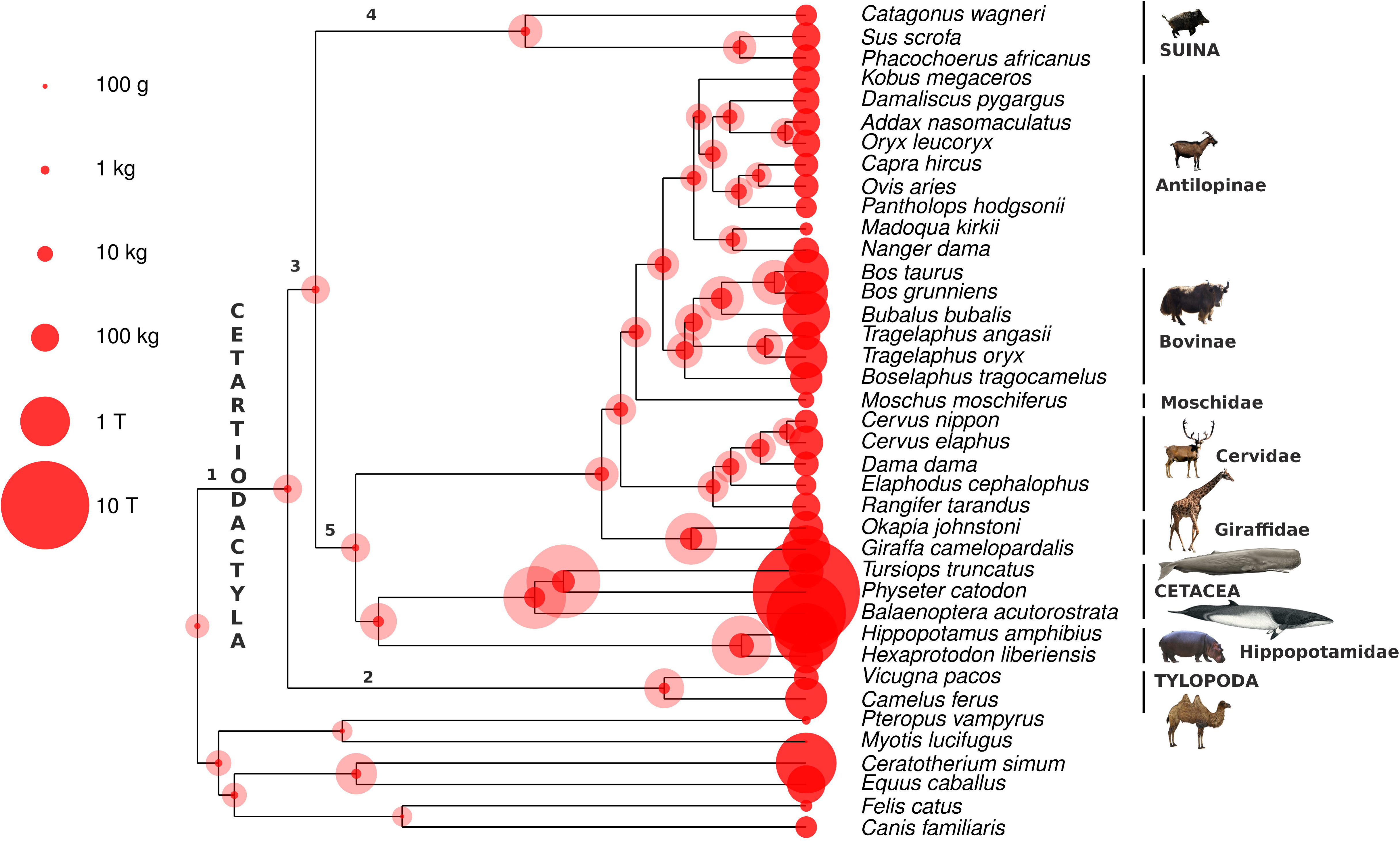
Reconstruction of body mass in the Cetartiodactyla using the integrated approach. All three LHTs were included in the analysis as well as both dS and dN/dS as dependent variables. Dark- and light-shaded disks indicate 95% credibility intervals for ancestral body masses (in grams). An uniform prior distribution was used to calibrate divergence times; see electronic supplementary material, figure S1. Numbers are labels for internal branches.

The estimated divergence times estimated by coevol on the reduced data set appeared to be robust to the choice of the prior on divergence times (uniform versus birth death) (see electronic supplementary material, figure S1). In particular, within the Cetartiodactyla clade, estimated dates differ by no more than 2 Myr between the analyses under the two alternative priors.

### dN/dS vs. life-history traits correlations in the Cetartiodactyla

Using a data set of 3,022 genes and 3.4 millions of positions, we estimated dN/dS in terminal branches of the tree and correlated it to three life-history traits. Significant correlations were detected between dN/dS and body mass (r = 0.56 P < 1.10^−3^, figure 2a), longevity (r = 0.59, P < 1.10^−3^) and sexual maturity (r = 0.56 P < 1.10^−3^), thus confirming in the Cetartiodactyla the influence of these traits on coding sequence evolutionary processes. However, no significant correlation was found between Kr/Kc (ie, ratio of radical over conservative amino-acid substitution rate) and any of the three life-history traits.

**Figure 2.**
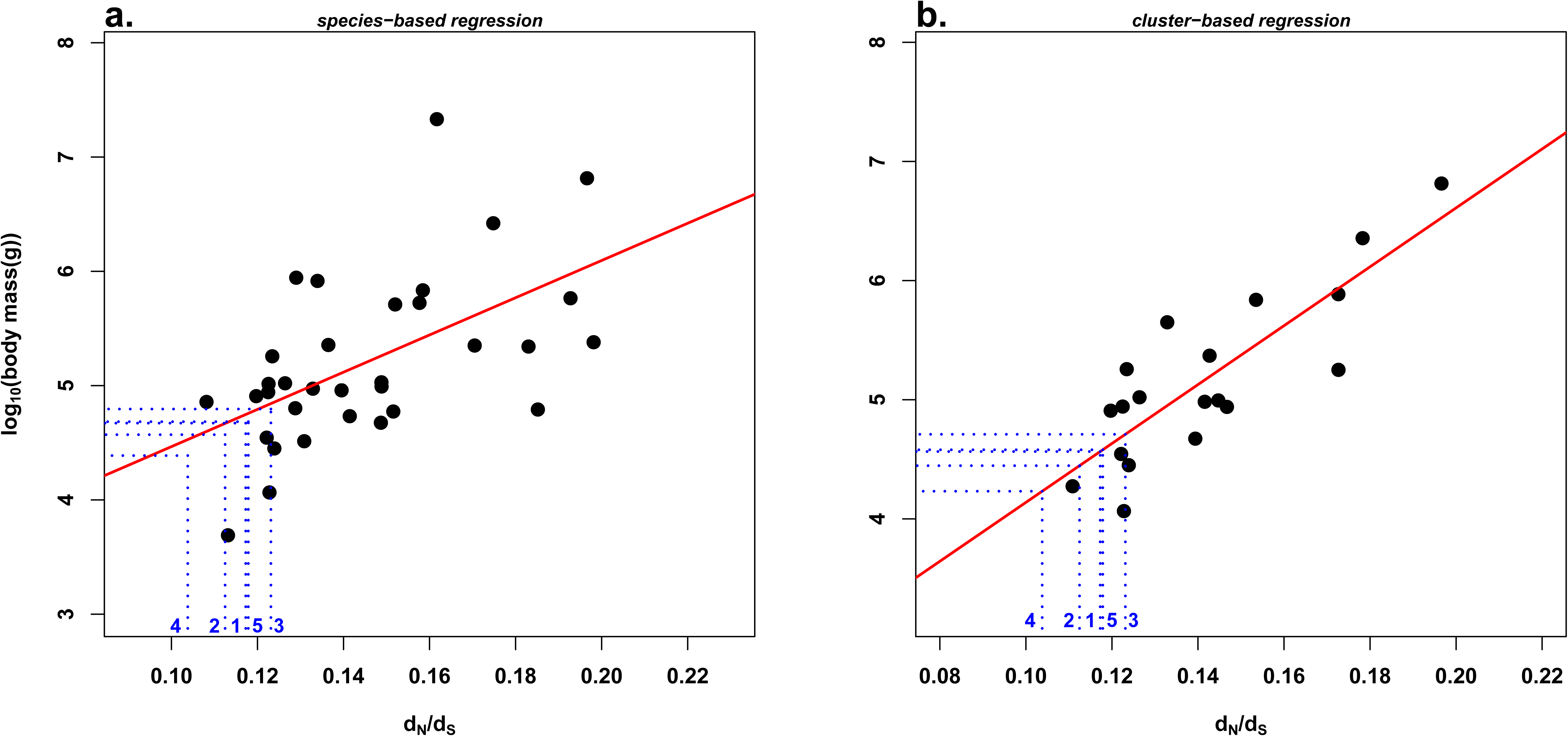
Relationship between log_10_(body mass) and dN/dS in Cetartiodactyla. **a.** Each dot represents a species. dN/dS values were obtained using substitution mapping. The regression line of log_10_(body mass) on terminal branch dN/dS is shown in red. The blue dashed lines show ancestral body mass predictions for five basal branches of Cetartiodactyla. Branch labels: see figure 1. **b.** Each dot represents a cluster of monophyletic species. Body mass value of a cluster corresponds to the mean of the log10-transformed values of its representatives.

Because current life-history traits might not accurately predict the long-term effective population size, we constructed clusters of monophyletic species by combining the substitution information across all its terminal branches. This procedure led to stronger correlations for body mass (r = 0.83, P < 1.10^−4^, figure 2b), longevity (r = 0.78, P < 1.10^−4^), and age of sexual maturity (r = 0.72, P < 1.10^−3^) when LHTs were averaged across our sampled species or across all species available in the AnAge database (body mass: r = 0.82, P < 1.10^−4^; longevity: r = 0.81, P < 1.10^−4^ ; age of sexual maturity: r = 0.85, P < 1.10^−5^).

We estimated ancestral dN/dS values in five basal branches of the Cetartiodactyla tree (figure 1) via substitution mapping. These fell in the lowest range of extant species dN/dS, and are therefore each suggestive of relatively small body-mass – 10-100kg – in the corresponding ancestor (blue dotted lines in figure 2). The strength and slope of the dN/dS - LHTs correlations and the estimated ancestral dN/dS values were not sensitive to the use of alternative topologies in which the branching order among outgroups or the position of the Nile lechwe differed (see electronic supplementary material, Appendix B). Using the reduced data set of 1,223 genes with less missing data also returned similar results (see electronic supplementary material, table S5).

### Modelling the correlated evolution between substitution parameters and LHTs

In order to properly account for the covariation between LHTs and substitution rates, we used the coevol program on the reduced data set, which models the evolution of molecular and life-history variables as a multivariate Brownian motion. The analysis under coevol indicates a negative correlation of the synonymous substitution rate dS, and a positive correlation of dN/dS, with the three LHTs (table 1, correlation coefficients vary by no more than 0.03 between both chains). Concerning dS, the correlation is essentially driven by sexual maturity (r = −0.61), which suggests a generation-time effect on the mutation rate. In contrast, for dN/dS, the correlation is primarily with body-mass (r = 0.52). Overall, the correlation estimated by this integrative modeling strategy are consistent with the mapping-based methods (see above).

**Table 1.** Relationships between life-history traits and molecular substitution rates in the Cetartiodactyla using the integrated approach. Values are given for chain 1.

|  | body mass | longevity | sexual maturity |
| --- | --- | --- | --- |
| dS | -0.56** | -0.53** | -0.61** |
| dN/dS | 0.52** | 0.44* | 0.23 |

\*\* $p > 0.975$ or $< 0.025$ , \* $p > 0.95$ or $< 0.05$

The body mass inferred by coevol in the early lineages of the group suggest a relatively small ancestor (95% credible interval between 500g to 115kg). This should be contrasted with what would be obtained without any information coming from molecular sequences – thus, based on a univariate log-Brownian model for body mass (credible interval between 10kg and 770kg).

Analyses using either dS or dN/dS as the single dependent variable against all three LHTs suggest that the correlation between LHTs and dS is the primary driver of the inference of smaller ancestral body mass, although dN/dS also appears to contribute a weak signal in favor of a small-bodied ancestor (table 2).

**Table 2.** Estimated body mass (95% credibility interval) at the root of the Cetartiodactyla. All *coevol* and the *ancov* analysis, that does not use molecular data, are shown.

|  | dS, dN/dS - traits | dS - traits | dN/dS - traits | traits |
| --- | --- | --- | --- | --- |
| chain 1 | 530 g - 89 kg | 570 g - 97 kg | 7.6 kg - 498 kg | 12 kg - 760 kg |
| chain 2 | 640 g - 115 kg | 590 g - 111 kg | 8.5 kg - 607 kg | 10 kg - 770 kg |

### GC3-based ancestral reconstruction

We analyzed the variation of GC3, a variable known to be influenced by LHT in mammals, to approach ancestral life-history traits. We retrieved strong correlations in the reduced data set between the across-genes mean GC3 and LHT in Cetartiodactyls despite the low range of observed mean GC3 (49.5%−51.5%): r = −0.65, P < 1.10^−4^ for body mass, r = −0.69, P < 1.10^−5^ for maximum longevity and r = −0.67, P < 1.10^−4^ for age of sexual maturity (see electronic supplementary material, figure S3), in agreement with a more pronounced action of gBGC in small-sized cetartiodactyls presumably having larger effective population sizes (Romiguier et al. 2010; Lartillot 2013).

GC3 cannot be directly used to estimate LHTs because of phylogenetic inertia. Following Romiguier et al. (2013), we identified eleven pairs of closely related species and calculated for each the correlation coefficient of GC3 across genes, and its divergence time-corrected version, γ. We observed a strong correlation between body mass and γ across the eleven pairs of extant species (τ = 0.68, P < 0.02, figure 3). Then we calculated γ in five pairs of ancestral Cetartiodactyl species (figure 1), ancestral GC3 being estimated based on a non-homogeneous, non-stationary model of sequence evolution. Similarly to the dN/dS analysis, we observed that pairs of basal Cetartiodactyl species all exhibited a low level of GC3 conservation, again suggesting a relatively small body size for these ancestors. Similar results were obtained when divergence times were taken from our coevol analysis rather than TimeTree (see electronic supplementary material, figure S4), the correlation between body mass and γ being slightly stronger in this analysis (τ = 0.79, P < 0.004). Two of the five ancestral branches were suggestive of medium-size ancestors, not small-size ancestors. These are the Tylopoda and Suina ancestral branches, each corresponding to a period of time longer than 25My.

**Figure 3.**
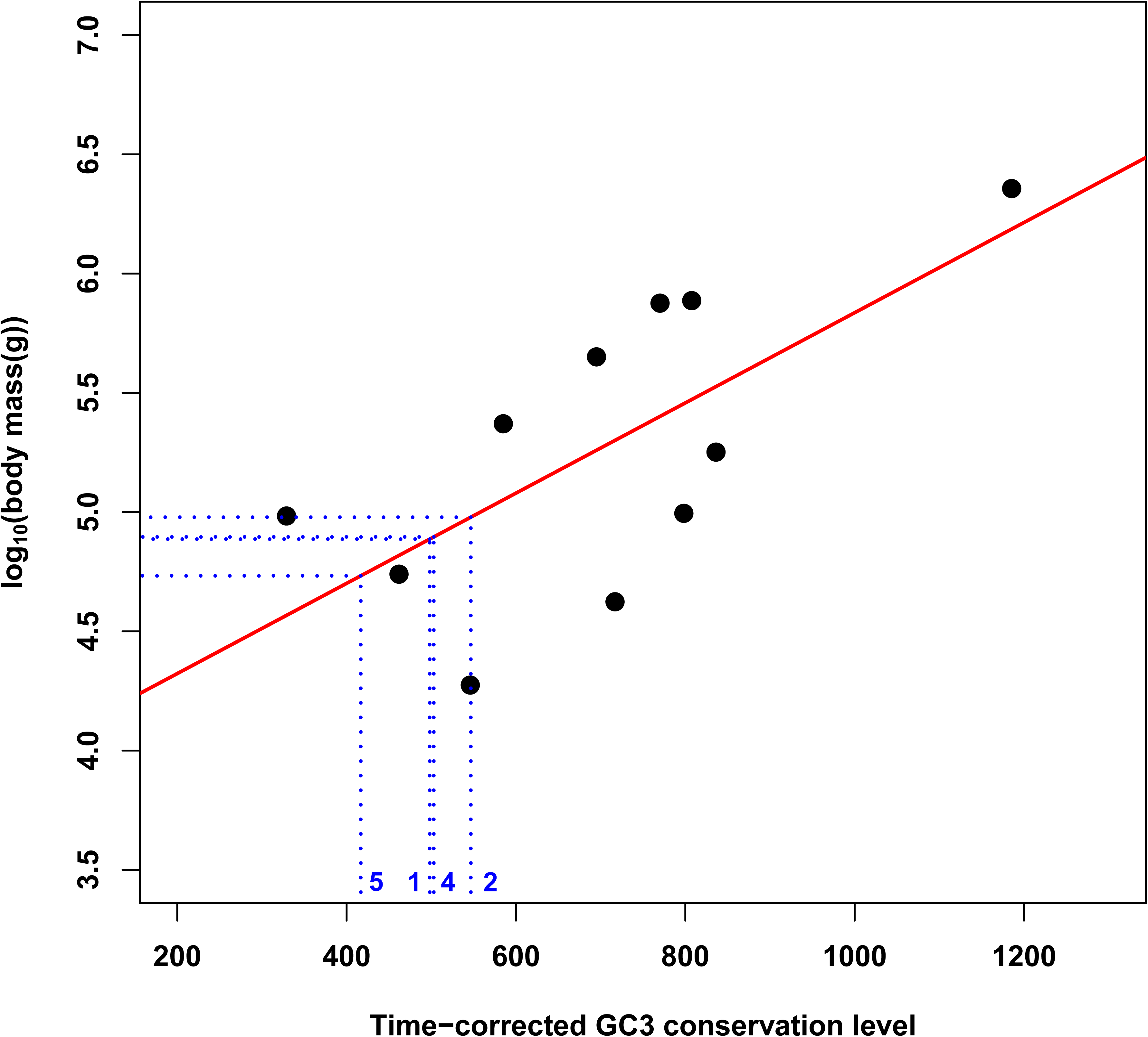
Relationship between log_10_(body mass) and time-corrected GC3 conservation level. Each dot represents a pair of cetartiodactyl species. The x axis corresponds to γ=-t/log(τ), where t is the divergence time taken from TimeTree and τ the the Kendall rank correlation coefficient. The regression line of log_10_(body mass) on time-corrected GC3 conservation level is shown in red. The blue dashed lines show the ancestral body mass predictions for four of the five most basal branches of Cetartiodactyla (branch 3 is not displayed because of its particularly low γ value: 98). Branch labels: see figure 1.

## DISCUSSION

Our goal in this study was to test the ability of molecular-aided method for ancestral trait reconstruction in a difficult case study – the Cetartiodactyla. We show that between lineages patterns of variation in synonymous and non-synonymous substitution rates and in GCcontent all point towards relatively small ancestors for this group. Our analyses indicate that the per year synonymous substitution rate, the dN/dS ratio and GC-content dynamics all carry some signal regarding ancestral traits in cetartiodactyls. In terms of the underlying molecular evolutionary mechanisms, this signal is possibly mediated by two distinct effects, on generation-time (for the variation in absolute synonymous substitution rate) and on effective population size (for the variation in dN/dS and GC content), both of which are known to correlate with body-mass in vertebrates (Bromham 2002; Woolfit and Bromham 2005; Nabholz et al. 2008). Thus, in at least two respects, molecular evolution in ancestral cetartiodactyls was similar to molecular evolution in currently small-sized taxa and different from molecular evolution in currently large-sized taxa.

The inference of a small cetartiodactyl ancestor is in agreement with the fossil record – and this, even though the vast majority of current species are large in size. Molecular-aided point estimates of ancestral body mass are about one order of magnitude below those of DNA-unaware methods (between 0.5 and 115 kg with molecular information, versus 10kg to 770kg without this information). This illustrates a point already observed in another phylogenetic context (evolution of the optimal growth temperature in Archaea, (Groussin and Gouy 2011; Horvilleur and Lartillot 2014), namely, that molecular information makes it possible to infer global trends in trait evolution along a tree, which classical comparative approaches are not able to identify, at least without fossil information. Our analysis therefore illustrates the potential of molecular data to solve difficult problems regarding ancestral state reconstruction, in situations where classical comparative methods have inherent limitations.

The most ancient fossils of Cetartiodactyla are found in the Eocene, i.e., at least a ten million years after the origin of the group according to molecular dating (Murphy et al. 2007; Meredith et al. 2011, this study). Fossils such as *Diacodexis* are difficult to pinpoint precisely onto the phylogeny of extant taxa, and can only be used as a raw guide for comparison with our estimated body masses in basal branches. Still, our reconstruction suggests that cetartiodactyl ancestors have remained small-sized for an extended period, from the Paleocene to the end of the Eocene, during which paleontology indeed only witnesses small-sized species. We infer independent evolution toward larger sizes in the major Cetartiodactyla lineages (figure 1), namely Tylopoda, Suina, Ruminantia and Cetacea. This is again consistent with the fossil record of these groups, particularly Tylopoda, in which families such as *Oromerycidae* consisted of small-sized species, and the well documented Cetacea (Montgomery et al. 2013).

Our approach to rate analysis relies on a single, supposedly known species tree and concatenated sequences. It was recently suggested that such an approach could be biased in case of incomplete lineage sorting, when gene trees differ from the species tree (Hahn & Nakhleh 2016, Mendes et al. 2016). Incomplete lineage sorting, however, can only be effective if internal branch length is of the order of magnitude of, or lower than, within species polymorphism (Scornavacca and Galtier 2017). Published population genomic data in Cetartiodactyla indicate that coding sequence polymorphism is of the order of 0.001-0.1% (Wu et al. 2014, Yim et al. 2014, Kardos et al. 2015, Choi et al. 2015).

Only two internal branches in our reference RaxML tree are that short (see electronic supplementary material, figure S2, appendix B), which represent a tiny fraction of the whole tree length. We conclude, following Scornavacca and Galtier (2017), that incomplete lineage sorting is probably a negligible issue in this analysis. Please also note that our results are robust to changes in the reference phylogeny (Table S5).

Interestingly, the Bayesian analysis conducted here is mainly driven by dS, with dN/dS contributing only a minor amount of information, whereas substitution mapping suggests quite a strong link between dN/dS and LHTs. The reason for such a discrepancy between the two approaches is yet unclear. Please note that a time tree is used in the Bayesian analysis, whereas dN/dS were estimated using a non-ultrametric tree with substitution mapping. This could explain in part the discrepancies between the two analyses. In addition, clustering species by clade and analysing cluster averages yielded a substantial increase in correlation coefficient, compared to the basic analysis (figure 2). This would not be expected if the logarithm of rates and traits were evolving according to a multivariate Brownian diffusion process, as assumed by the coevol model, and suggests that not all the information contained in the data was captured by our analysis. Modelling the correlated evolution of molecular and life-history traits is a complex, computationally demanding task, but a necessary one if one wants to properly account for phylogenetic inertia (Lartillot and Poujol 2011). This study confirms the potential of an integrated, Bayesian approach while offering suggestions for improvement of existing models. Besides these methodological aspects, it should be noted that the correlation between rates and traits is probably imperfect in nature – an intrinsic limitation to our approach, which probably determines in the first place the width of credibility intervals.

The success of DNA-aided methods in reconstructing small cetartiodactyl ancestors, such as independently indicated by fossil evidence gives credit to other analyses for which fossil data are lacking — in particular, the suggestion by Lartillot and Delsuc (2012) and Romiguier et al. (2013) of a relatively large and long-lived placental ancestor. Given the controversy on the age of this ancestor and the uncertain interpretation of mammalian fossils from the Cretaceous (Ronquist et al. 2016; Springer et al. 2017), relying on the correlations between life history traits and substitution processes is probably the optimal way to currently approach ancestral states in mammals. Time-independent molecular variables, such as the dN/dS ratio, are particularly appropriate when dates are controversial, as in the case of early placental evolution. Another promising approach involves analysing the variance, not just the mean, of substitution rates across genes (Wu et al. 2017).

Our approach, however, is only applicable when strong correlations exist between rates and traits, which is the case in some but not in every taxonomic group (Bromham 2002; Lourenço et al. 2013; Figuet et al. 2016; Rolland et al. 2016). Such correlations exist because life-history traits influence substitution rates through the mutation rate, effective population size, recombination rate, or other evolutionary forces. Further understanding the link between rates and traits, e.g. via population genomic studies, should enhance our ability to model these relationships and improve ancestral reconstructions.

## Acknowledgements

The authors are grateful to the Lunaret zoo, Servion zoo, Pescheray zoo, Gramat zoo, Cédric Libert, Elodie Trunet, Eugène Chabloz, Jean-Marc Charpentier, and Pierre Delrieu for their great help with sampling. They are particularly grateful to the San Diego Frozen Zoo and Cynthia Steiner for providing RNA data from species that were particularly difficult to sample. They also thank Laurent Duret and Jonathan Romiguier for helpful discussion and the Montpellier Bioinformatics & Biodiversity platform for support regarding computational aspects. This work was supported by European Research Council advanced grant 232971 (PopPhyl) and Agence Nationale de la Recherche grant ANR-10-BINF-01-01 (Ancestrome).

